# Influenza A Virus Exacerbates Group A Streptococcus Infection and Thwarts Anti-bacterial Inflammatory Responses in Murine Macrophages

**DOI:** 10.1101/2022.03.10.483890

**Authors:** Johann Aleith, Maria Brendel, Erik Weipert, Michael Müller, Daniel Schultz, Ko-Infekt Study Group, Brigitte Müller-Hilke

## Abstract

Seasonal influenza epidemics pose a considerable hazard for global health. In the past decades, accumulating evidence revealed that influenza A virus (IAV) renders the host vulnerable to bacterial superinfections which in turn are a major cause for morbidity and mortality. However, whether the impact of influenza on anti-bacterial innate immunity is restricted to the vicinity of the lung or systemically extends to remote sites is underexplored. We therefore sought to investigate intranasal infection of adult C57BL/6J mice with IAV H1N1 in combination with bacteremia elicited by intravenous application of Group A Streptococcus (GAS). Co-infection *in vivo* was supplemented *in vitro* by challenging murine bone marrow derived macrophages and exploring gene expression and cytokine secretion. Our results show that viral infection of mice caused mild disease and induced the depletion of CCL2 in the periphery. Influenza preceding GAS infection promoted the occurrence of paw edemas and was accompanied by exacerbated disease scores. *In vitro* co-infection of macrophages led to significantly elevated expression of TLR2 and CD80 compared to bacterial mono-infection, whereas CD163 and CD206 were downregulated. The GAS-inducible upregulation of inflammatory genes, such as *Nos2*, as well as the secretion of TNFα and IL-1β were notably reduced or even abrogated following co-infection. Our results indicate that IAV primes an innate immune layout that is inadequately equipped for bacterial clearance.

## 1 Introduction

Seasonal influenza is a major cause of respiratory disease that affects 5 – 10% of the global population annually with an estimated death toll of up to 500,000 [1,2]. The segmented genome of influenza A virus (IAV) combined with an error-prone RNA polymerase enables the periodical emergence of new strains with elevated pandemic capacities, which annually challenge humankind yet devoid of adequate adaptive immunity [3,4]. The most prominent paradigm for the dramatic consequences of an influenza pandemic is the 1918/1919 flu that caused roughly 50 Million casualties [5]. Notably, the vast majority of fatal cases were attributed to secondary bacterial infections predominantly caused by pneumococci and hemolytic streptococci [6,7]. Along these lines, excess morbidity due to bacterial superinfection with the nasopharyngeal colonizers *S. pneumoniae, S. aureus* and *S. pyogenes* (Group A Streptococcus, GAS) was confirmed for the most recent influenza pandemic in 2009 [8]. As of yet, there is neither a licensed vaccine against S. *aureus* nor against *S. pyogenes* that would help contain invasive infections with these pathogens during future influenza pandemics [9–11].

Several modes by which an immune response against IAV supports viral clearance yet fails to oppose bacterial pathogens have been suggested [1]. For instance, Okamoto and colleagues demonstrated that IAV infection led to the presentation of hemagglutinin (HA) by epithelial cells, which is utilized by GAS to breach cellular barriers [12,13]. Other groups reported that HA, among other viral proteins, caused the exposure of receptors that act as adhesins for bacterial attachment and invasion [14–16]. Others showed that viral infection caused damage of the respiratory epithelium, expediting initial bacterial adherence [6,17,18]. Moreover, experimental data indicated that IAV paves the way for the dissemination of opportunistic bacterial pathogens by impacting the innate immune response, which is critical for bacterial containment [19,20]. In fact, the virus was shown to induce an increased secretion of anti-inflammatory interleukin (IL-)10 as well as inflammatory type I and type II interferons (IFNs), which was associated with both, impaired phagocytic activity by pulmonary immune cells and diminished production of chemokines [14,19,21–24].

Together, these data illustrate some aspects of post-influenza pneumonia and the interplay of viral and bacterial pneumopathogens in life-threatening infections. While the aforementioned studies focused on bacterial superinfections of the respiratory tract, we were intrigued that seasonal or pandemic influenza outbreaks seem to coincide with a broad spectrum of invasive GAS-associated infectious diseases like necrotizing fasciitis, pneumonia and bacteremia [8,25–30]. We therefore asked whether pulmonary IAV can also alter systemic innate immunity and facilitate secondary bacterial insults at remote sites. We were particularly interested in the impact IAV exerts on the response of macrophages – immune cells that are indispensable for initial anti-streptococcal resistance [19,31,32]. We established co-infection models that *in vivo* combined respiratory IAV infection with GAS bacteremia and *in vitro* investigated primary macrophages for their potential to respond to both pathogens simultaneously.

## 2 Materials and Methods

### 2.1 Pathogens

Pandemic influenza A virus (IAV) A/Germany-BY/74/2009 (H1N1pdm09) propagation and titer determination was performed as previously described [33]. In brief, IAV was replicated in Mardin-Darby canine kidney II (MDCKII) cells using minimal essential medium supplemented with 0.2% bovine serum albumin and 2 μg/mL N-Tosyl-L-phenylalanin-chlormethylketon (Sigma). For the determination of the tissue culture infectious dose 50 (TCID_50_), virus suspensions were serially diluted and applied to MDCKII cultures. Cells were then incubated for three days at 37°C and 5% CO_2_ followed by examination of cytopathogenicity.

*Streptococcus pyogenes* (Group A Streptococcus, GAS) strain AP1 of the *emm1* (M1) serotype was originally acquired from the World Health Organization Collaborating Center for Reference and Research on Streptococci (Prague, Czech Republic). Bacteria were thawed onto Colombia agar plates containing 5% sheep blood (Becton Dickinson) and were cultured overnight followed by storage at 4°C for up to three weeks. Colonies were picked from the plate, suspended into Todd-Hewitt broth (THB, Becton Dickinson) and cultured overnight at 37°C and 5% CO_2_. The suspension was diluted 20-fold in THB and bacteria were incubated until exponential phase of growth was reached. Subsequently, bacteria where washed thrice with PBS (Thermo Fisher) prior to their application in mice and *in vitro* infection models, respectively. The determination of colony forming units (CFU) was performed the following day by counting of serially diluted suspensions.

### 2.2 Animals

C57BL/6J mice were initially purchased from Charles River. Mice were bred in the animal core facility under specific germ-free conditions. Animals were transferred to individually ventilated cages prior to infection experiments and were housed at a 12-hour light/dark cycle, an ambient temperature of 22 ± 2°C and 50 ± 20% humidity. Food and water were provided *ad libitum*. Animal experiments were reviewed and approved by the ethics committee of the State Department for Agriculture, Food Safety and Fishery in Mecklenburg-Western Pomerania under the file reference number 7221.3-1-017/19.

### 2.3 In vivo infection models and clinical scoring

For the induction of viral infections, 20 μL of a suspension containing 1.5 × 10^5^ TCID_50_ IAV were applied to both nostrils of 20- to 22-week old male mice under anesthesia by isoflurane inhalation. This volume was chosen in order to guarantee an infection of both, the upper and lower respiratory tracts [34]. Applying the same volume of PBS only served as the negative (healthy) control. Mice were subsequently monitored daily for 16 days for alterations in body weight relative to the day of infection (day 0). On days 2, 4 and 7, a maximum of 80 μL of anti-coagulated blood was drawn by saphenous venipuncture using a 25G needle followed by centrifugation and collection of plasma. On day 16, mice were anesthetized with 75 mg of Ketamine (Pharmanovo) and 5 mg Xylazin (Bayer) per kg bodyweight. Subsequently, mice were exsanguinated by cardiac puncture. Mice were then sacrificed by cervical dislocation and lungs were excised, snap frozen and stored at -80°C for later analyses.

In order to induce bacteremia, GAS was diluted in PBS and 1 × 10^5^ CFU were applied in a 100 μL volume by injection into the lateral tail vein. Intravenous injection of PBS served as a control. For co-infection, IAV was applied as described above either two days prior or subsequent to bacterial infection. Mice were given tramadol (Ratiopharm) in drinking water for analgesia. Animals were monitored following bacterial infection for a maximum of 14 days or until humane endpoints were reached. Sepsis severity was assessed by a scoring system that incorporated the assessment of macroscopic signs of burden as previously described [35,36]. In brief, scores of four categories were added together to provide an estimate for overall sepsis activity: i) weight loss of ≥ 5% (Score 5), ≥ 10% (Score 10), ≥ 20% (Score 20, humane endpoint); ii) appearance deviations such as piloerection (Score 5), high myotonicity or scruffy orifices (Score 10), convulsions or paralysis (Score 20, humane endpoint); iii) impairment of consciousness such as suppressed activity or limited reaction to stimuli (Score 5), self-isolation or lethargy (Score 10), perpetual pain vocalization or apathy (Score 20, humane endpoint) and iv) signs of impaired respiratory quality or inflammation such as edemas on small body areas (Score 5), disseminated edemas or labored breathing (Score 10), open wounds or gasping (Score 20, humane endpoint).

Mice were sacrificed as described above upon reaching the end of the observation period, at any humane endpoint or when reaching an overall sepsis score of ≥ 20. Cardiac blood samples were plated on blood agar and medial arthrotomy on both knee joints was performed under a stereo microscope followed by plating of the synovial fluid on blood agar. Agar plates were subsequently incubated overnight and examined for the presence of β-hemolytic bacteria. Hind paws were extracted, snap frozen and stored at -80°C for the analysis of eicosanoids.

### 2.4 Eicosanoid extraction and analysis

Lipidomics analyses were performed as previously described [35]. In brief, paw samples were chilled in liquid nitrogen, pulverized and 50 mg of the resulting powder was immersed in 500 μL cold methanol containing 0.1% butylated hydroxytoluene and 500 μL ice cold water. 100 μL deuterated internal standards containing 12-HETE-d_8_, 13-HODE-d_4_, PGE_2_-d_4_ and Resolvin D1-d_5_ (each 100 ng/mL, Cayman Chemicals) were subsequently added followed by an additional lysis step with matrix B at 6 m/s for 45 s on a FastPrep (MP Biomedicals). Following this, 300 μL sodium acetate (1 M) was added on ice and 10 M acetic acid was added until pH 6 was reached. Solid phase extraction was performed on methanol and sodium acetate conditioned Bond Elut Certify II cartridges (Agilent). After loading the samples, cartridges were washed with 50% methanol. Elution of eicosanoids was carried out by addition of hexane/ethyl acetate (75/25) containing 1% acetic acid.

For measurements, paw extracts were dried under nitrogen flow using a TurboVap (Biotage) and reconstituted in 70 μL 25% acetonitrile. Separation was done on a Gemini NX-C18 column (3 μm, 100 × 2 mm) utilizing an Agilent 1200 series HPLC systems. Dynamics multiple reaction monitoring MS/MS was executed using a 6460 series triple quadrupole tandem mass spectrometer (Agilent) with electrospray ionization in negative mode. Calibration by internal and external standards was performed as previously described [35]. Agilent Mass Hunter Qualitative Analysis software and Agilent Mass Hunter Quantitative Analysis software (both version B.07.00) were used for MS data analysis. Quantities of individual eicosanoids were standardized to a mean of 0 and a standard deviation of 1 for data visualization.

### 2.5 Isolation of RNA and DNA from lung samples

Lung samples were submerged in liquid nitrogen, slightly fragmented and weighed. Sixty to one hundred twenty milligrams were transferred to lysis tubes containing bashing beads (Zymo Research) and 1 mL TRIzol (Thermo Fisher). Lung fragments were subsequently homogenized at 4,000 rpm for 4 × 20 s using a FastPrep. Samples were then centrifuged at 10,000 × *g* for 7 min at 4°C and transferred into new tubes. Apart from centrifugation at 4°C, the following steps were conducted at room temperature. After resting for 5 min, 200 μL chloroform (Sigma) was added and samples were extracted for 3 min. Subsequently, samples were centrifuged for 15 min at 12,000 × *g*. The RNA-enriched upper phase was mixed with 500 μL 2-propanol, incubated for 10 min and centrifuged at 12,000 × *g* for 10 min. RNA pellets were suspended in 75% Ethanol followed by centrifugation at 7,500 × *g* for 5 min. Supernatants were subsequently discarded, pellets were dried and dissolved in 40 μL RNAse-free water by incubation at 60°C for 15 min. RNA contents were then determined photometrically on a NanoDrop (Thermo Fisher). DNA was isolated by precipitation of the appropriate phase upon addition of 300 μL ethanol, incubation for 3 min and centrifugation for 5 min at 2,000 × *g*. The resulting pellet was then washed twice by 30 min incubation with 0.1 M sodium citrate (pH 8.5) in 10% ethanol. DNA samples were subsequently suspended in 75% ethanol and incubated for 20 min. After centrifugation, supernatants were discarded, pellets were dried and dissolved by incubation in 8 mM NaOH for 10 min. DNA contents were determined fluorometrically using the Qubit 1X dsDNA Assay Kit to the manufacturer’s instructions (Thermo Fisher).

### 2.6 Lung pathogen burden and gene expression

Primer pairs were designed for the detection of IAV H1N1 matrix protein, nucleoprotein and hemagglutinin in murine lung extracts according to the strain specific sequences found at https://www.fludb.org/brc/fluStrainDetails.spg?strainName=A%2FGermany-BY%2F74%2F2009%28H1N1%29&decorator=influenza (supplementary Table I). For this, RNA was isolated as described above and 500 ng were reverse transcribed using the High Capacity cDNA Reverse Transcription Kit (Thermo Fisher) according to the manufacturer’s instructions. Twenty-five nanograms of the resulting cDNA together with 500 nM of the primer pairs were submitted to qPCR using the PowerUP SYBR Green Mastermix (Thermo Fisher). The amplification reaction was monitored on the ViiA 7 Real-Time PCR System running on the QuantStudio Real Time PCR Software V1.3 (Thermo Fisher). The size of the respective amplicons was confirmed by 2% agarose gel and ethidium bromide staining. Primer pairs for the detection of GAS strain AP1 specific genomic DNA were designed according to sequence information found at https://www.ncbi.nlm.nih.gov/nuccore/CP007537?report=genbank (supplementary Table II). A total of 20 ng DNA from lung extracts were used together with 500 nM of the primer pairs for qPCR as described above, followed by confirmation of amplicon sizes on agarose gels. Gene expression analyses were performed on 25 ng cDNA that was obtained from reverse transcribed lung RNA. For qPCR analysis, TaqMan primer pairs and probes (Thremo Fisher) were used for *Ccl2* (assay ID: Mm00441242_m1) and *Ifnb1* (Mm00439552_s1) utilizing *Gapdh* (Mm05724508_g1) as a reference gene. All reactions were amplified using the TaqMan Gene Expression Master Mix (Thermo Fisher).

### 2.7 Bone marrow derived macrophage infection model

C57BL/6J mice used for bone marrow isolation had a median age of 10 weeks (range 7 – 42 weeks) and 30% were female. Bone marrow was obtained from long bones by centrifugation as previously described [37]. The resulting pellet was subsequently suspended in Dulbecco’s Modified Eagle’s Medium (DMEM) supplemented with 10% fetal calf serum (FCS), 5 IE/mL Penicillin, 5 μg/mL Streptomycin, 2 mM L-Glutamine (Thermo Fisher), 10 mM HEPES and 1 mM sodium pyruvate (PAN Biotech). After determination of vital cells using a hemocytometer and trypan blue (Thermo Fisher), cells were seeded into 6-well culture plates (Greiner) at a density of 3 × 10^5^ cells per cm^2^ in 2 – 5 mL supplemented DMEM. The differentiation to macrophages was initiated at day 0 by the addition of 20 ng/mL macrophage colony-stimulating factor (M-CSF, R&D Systems). Cells were cultured afterwards at 37°C and 5% CO_2_ for 7 days including the replacement of supplemented DMEM and replenishment of M-CSF at days 1 and 4. For viral infection (day 7, t_0_), supplemented DMEM was refreshed and 4 × 10^5^ TCID_50_ IAV were added. Following this, infected or uninfected macrophages were incubated for 48 h upon which the cells were either collected for downstream analyses or submitted to bacterial (super-)infection (day 9). In case of the latter, supplemented DMEM was removed, the cells were washed thrice with PBS and Minimal Essential Medium α containing additional nucleosides and 10% FCS (Thermo Fisher) was added. GAS was then applied at 4.5 × 10^6^ CFU. Subsequently, macrophages were incubated for 6 h followed by sample collection.

### 2.8 Single cell analysis by flow cytometry

Gentle detachment of macrophages from culture plates was carried out by washing with PBS and subsequently incubating with 5 mL PBS containing 10 mM EDTA for 10 min. Culture plates were tapped multiple times and suspensions were collected afterwards. For increased yields, 0.7 mL accutase (Pan Biotech) was added for 10 – 15 min followed by alternately tapping and pipetting. Subsequently, another 0.7 mL accutase were added for an additional 10 – 15 min, tapping and pipetting were repeated and suspensions collected and pooled with the PBS/EDTA fraction. Finally, 1 mL supplemented DMEM was added and the remaining cells were obtained using a cell scraper (Sarstedt). Suspensions were centrifuged at 400 × *g* and 4°C for 5 min and cells were suspended in autoMACS Running Buffer (RB, Miltenyi Biotec) followed by counting. Antibody binding to CD16 and CD32 was prevented by incubation of macrophages with 0.5 μg Trustain FcX (Biolegend) in RB supplemented with 10% FCS for 10 min on ice. Subsequently, an antibody mixture containing 0.13 μg (anti-)F4/80:FITC (clone BM8), 0.5 μg CD163:APC (S150491), 0.25 μg CD206:BV605 (C068C2), 0.25 μg CD80:BV421 (16-10A1, Biolegend), 0.22 μg CD86:APC/Vio770 (PO3.3), 4.5 μL TLR2:PE (REA109) and 0.15 μg MHCII:PerCP/Vio770 (REA813, Miltenyi Biotec) was added and incubated for 20 min on ice in the dark. Cells were washed afterwards, suspended in RB and 7-Aminoactinomycin (7-AAD, Biolegend) was added at a concentration of 1.25 μg/mL for at least 5 min prior to measurement.

Data acquisition was performed on the Aurora spectral flow cytometer running on the SpectroFlo software v2.2.0.3 (Cytek Biosciences). Data analysis was conducted using the FlowJo software v10.7.1. Supplementary Fig. 1 illustrates the gating strategy. Live macrophages were identified as 7-AAD^−^F4/80^+^ singlets. This population was used for the subsequent determination of expression levels based on median fluorescence intensity (MFI) values and as a parent for measuring the proportions of subpopulations expressing different combination of the above-listed surface antigens. For dimension reduction, 10,000 macrophage events were down-sampled, concatenated and submitted to the algorithm t-distributed stochastic neighbor embedding (t-SNE) using an automated learning configuration (opt-SNE combined with the exact KNN algorithm and the Barnes-Hut gradient algorithm) with a perplexity of 50 and a maximum of 1000 iterations [38]. Unsupervised clustering of subpopulations expressing any combinations of the analyzed surface proteins was conducted by FlowSOM [39].

### 2.9 Macrophage gene expression

After aspirating cell culture supernatants, 700 μL of a chaotropic agent solution (Qiagen) was added to individual wells and cells were lysed by scraping and vigorous shaking. RNA was subsequently isolated using the RNeasy Plus Mini Kit (Qiagen) after the manufacturer’s instructions. Quantification of RNA contents were determined photometrically and 200 ng RNA was submitted to reverse transcription as described above. Amplification of cDNA was then performed by TaqMan Gene Expression Master Mix, primer pairs and probes for the relative quantification of *Ccl2, Cxcl2* (assay ID: Mm00436450_m1), *Ifnb1, Il1b* (Mm00434228_m1), *Il6* (Mm00446190_m1), *Il10* (Mm00439614_m1), *Mgl2* (Mm00460844_m1), *Nos2* (Mm00440502_m1), *Tgfb1* (Mm01178820_m1) and *Tnf* (Mm00443258_m1) using *Gapdh* as a reference gene. Polymerase chain reactions were performed on the Viia 7 System. In detail, samples were first incubated for 2 min at 50°C followed by 10 min at 95°C for polymerase activation. Subsequently, 40 automated cycles of PCR were performed that incorporated denaturation at 95°C for 15 sec and annealing and elongation at 60°C for 1 min. After each cycle the fluorescein amidite fluorescence signal was measured. Ct values were obtained when fluorescence intensities reached data-dependent and automatically defined thresholds. Quantification of gene expression was then performed by the 2^-ΔΔCt^ Method that incorporated normalization of the target gene Ct values to the reference gene (ΔCt) as well the difference between ΔCt values from uninfected and infected cells (ΔΔCt).

### 2.10 Cytokine analysis

Cytokine concentration in mouse plasma samples were quantified by a 3-plex LEGENDplex assay (Biolegend) that contained capture beads and detection antibodies for CCL2 (monocyte chemoattractant protein-1, MCP1), Interferon (IFN)γ and tumor necrosis factor (TNF)α. For the quantification of CCL2, Interleukin (IL-)1β, IL-6, IL-10 and TNFα in cell culture supernatants, a 5-plex LEGENDplex assay was used following the manufacturer’s guidelines. Data acquisition was performed on the Cytek Aurora flow cytometer. Cell culture supernatant concentrations of CXCL2 (macrophage inflammatory protein 2-α, MIP2-α) were determined by the CXCL2/MIP-2 DuoSet enzyme linked immunosorbent assay (ELISA) kit to the manufacturer’s instructions (R&D Systems). Horseradish peroxidase-catalyzed color reaction were initiated by addition of the TMB Substrate Kit (Biolegend) and quenched by 0.5 M sulfuric acid (Merck). The absorbance at 450 nm was measured on the Infinite M200 spectral photometer (Tecan).

### 2.11 Statistical analysis

Data analysis and visualization were performed using RStudio v1.2.5033 that ran R v3.5.1. Normalization was either performed by division of individual values from infection groups and control groups, respectively, or by feature scaling into a 0 – 1 range by the formula x_i_’ = (x_i_ – x_min_)/(x_max_ – x_min_). Heatmaps and hierarchical clusters were generated with the “pheatmap” package that incorporated feature scaling by standardization applying the formula 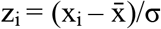. Calibration curves were fitted and samples values were estimated by n-parameter logistic regression using the “nplr” package. Two-sided statistical tests were used for the comparisons of group medians or means. Repeated measures (body weight trajectories) were compared by one-way and two-way analyses of variance (ANOVA), respectively. Probabilities of survival and incidences were compared by the logrank test. Bivariate interdependencies were evaluated by the Pearson product-moment correlation coefficient (r). Data sets were tested for normality by the Shapiro-Wilk test. Normal distribution of within-group raw or normalized variables was rejected when the test resulted in a p-value of < 0.05. Depending on the outcome of this test, univariate statistical analyses on variables that were normalized to respective controls were performed by the one-sample Wilcoxon signed rank test and the one-sample t-test, respectively. Two independent samples were compared with the Mann-Whitney U test or the t-test. Comparisons of variables between multiple groups were performed with the Dunn’s test and the Tukey HSD test, respectively, in combination with type I error correction using the Bonferroni-Holm method. A p-value of < 0.05 was considered statistically significant.

## 3 Results

### 3.1 Infection with influenza A virus H1N1 caused mild symptoms and induced a persistent immune reaction in the lung

In order to examine clinical manifestations of influenza, we used a model of intranasal infection with 2009 pandemic H1N1 IAV in adult mice (Fig. 1A). Intranasal application of PBS served as a control. Mice were monitored for relative weight loss post infection as a proxy for disease severity and indeed, exhibited minor reductions in body weight as early as two days after virus application (Fig. 1B). This trend continued until day seven after infection and resulted in a maximum weight loss of 5.5% ± 2.1% (mean ± SEM). Thereafter, body weight continuously increased and returned to starting values by day 14 suggesting robust recovery from infection. When comparing weight trajectories over the entire observation period using two-way analysis of variance (ANOVA), we found a statistically significant difference between infected mice and uninfected controls (p < 0.001). In accordance with the observed mild disease courses, we did not measure quantifiable amounts of the inflammatory cytokines TNFα and IFNγ in plasma samples from IAV infected animals (not shown). We did, however detect significant reductions of plasma CCL2 concentrations by 12.8% and 13.6% at days two and four after infection, respectively, relative to uninfected controls (Fig. 1C). By day seven, CCL2 plasma levels equalized between both groups (p = 0.65, not shown).

**Figure 1.**
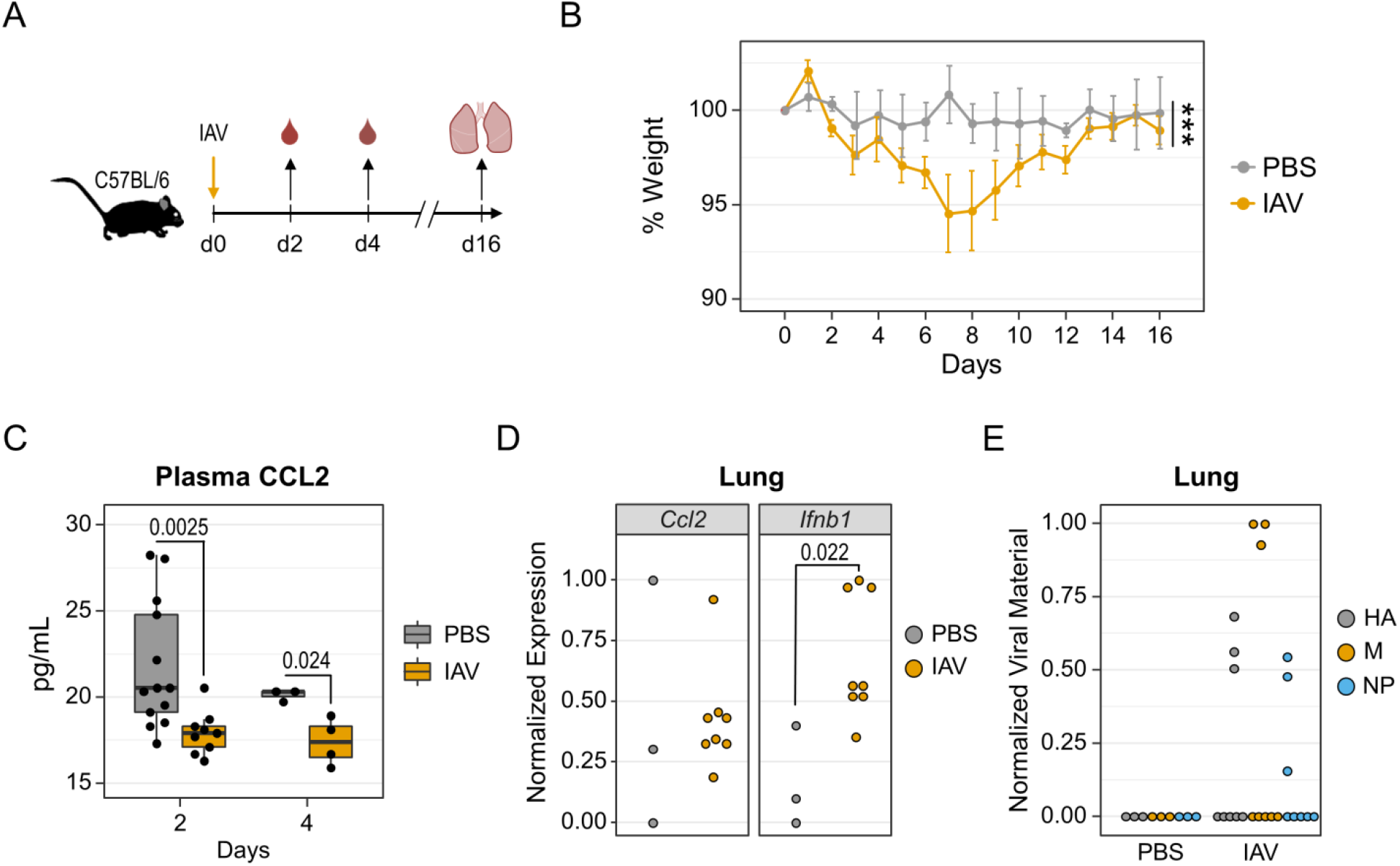
Influenza A virus infection induced minor weight loss and inhibited the production of CCL2. (**A**) Experimental Design. Mice were intranasally infected with influenza A virus (IAV, n = 8). PBS was adminstered as a control (n = 3). Blood samples were drawn on days 2 and 4 following infection. Lungs were excised at day 16. (**B**) Mean weight changes relative to day 0. Weight loss was confirmed by one-way ANOVA (p < 0.0001) in the IAV group and by two-way ANOVA (***p < 0.001) comparing the IAV group to PBS controls. Error bars depict the SEM. (**C**) Boxplots display CCL2 concentrations in plasma samples from uninfected controls (n = 13) and IAV infected mice (n = 9). The differences in samples sizes between days 2 and 4 are due to the fact that some animals were later subjected to the bacterial and co-infection experiments that are shown in Figure 2. Day 2 and Day 4 p-values result from Mann-Whitney U test and t-test, respectively. (**D**) Dotplots show normalized mRNA expression of *Ccl2* and *Ifnb1* in lung homogenates based on ΔCt values. p-value results from t-test. (**E**) Normalized viral loads based on Ct values for IAV specific genes in day 16 lung homogenates. HA: hemagglutinin, M: matrix protein, N: nucleoprotein.

In order to examine immune responses in the lower respiratory tract, we further performed gene expression analyses on whole lung homogenates that were obtained 16 days after IAV application. For this, we focused on mRNA expression levels of *Ccl2* and *Ifnb1*, as the former was altered in the periphery and the latter can be indicative of an anti-viral response. Protein data were not collected because of limited sample quantities. We found no meaningful differences in the expression of *Ccl2* between the IAV and control groups (Fig. 1D, left panel). Interestingly, *Ifnb1* expression was found to be significantly increased in lungs from infected mice (Fig. 1D, right panel). Given this prolonged upregulation of *Ifnb1*, we consequently utilized primer pairs for the detection of viral genes in lung samples that code for hemagglutinin, matrix protein and nucleoprotein (supplementary Fig. 2). We indeed detected IAV-specific RNA in 38% (3/8) of infected animals by quantitative PCR (Fig. 1E). False positive detection of unspecific targets was ruled out by confirming the expected amplicon melting temperatures (supplementary Fig. 3). However, the quantities of all three viral genes were generally low (C_t_ > 32) and might rather indicate residual viral antigen. Interestingly, two of the three samples that were positive for viral genes were also among the samples that expressed the highest amounts of *Infb1* indicating some correlation between both parameters.

In summary, we here show that an infection with IAV H1N1 in mice induced minor clinical manifestations that were accompanied by an early and reversible reduction of plasma CCL2 levels. Our data further show that barely detectable genetic material of the virus persisted in the lungs of some animals, which was accompanied by an ongoing type I IFN production.

### 3.2 Group A Streptococcal infection was aggravated following Influenza A virus infection

As CCL2 is integral to bacterial control [40–42], yet reduced during respiratory tract infection with IAV, we sought to investigate the clinical features of IAV superimposed bacteremia. To this end, we compared infection with bacteria only to co-infection models combining intranasal virus application with intravenous GAS infection in alternating succession (Fig. 2A). By monitoring for macroscopic symptoms following bacterial infection, we observed the occurrence of localized paw inflammation (Fig. 2B). Of note, the emergence of these edemas was accelerated and more frequent in post-influenza bacteremia (IAV+GAS) compared to bacterial infection only (GAS, p = 0.01) and pre-influenza bacteremia (GAS+IAV, p = 0.045), respectively (Fig. 2C). In detail, 80% (8/10) of mice in the IAV+GAS group exhibited signs of paw inflammation already one day after bacterial infection. In contrast, the incidence of paw edemas was increased to only 40% (4/10) in the GAS+IAV group as opposed to 20% (2/10) in the GAS only group, and this difference did not reach statistical significance (p = 0.33). We further analyzed eicosanoids from paw extracts and found that these immunologically active lipid metabolites were upregulated in some animals irrespective of the (co-)infection regimen (supplementary Fig. 4).

**Figure 2.**
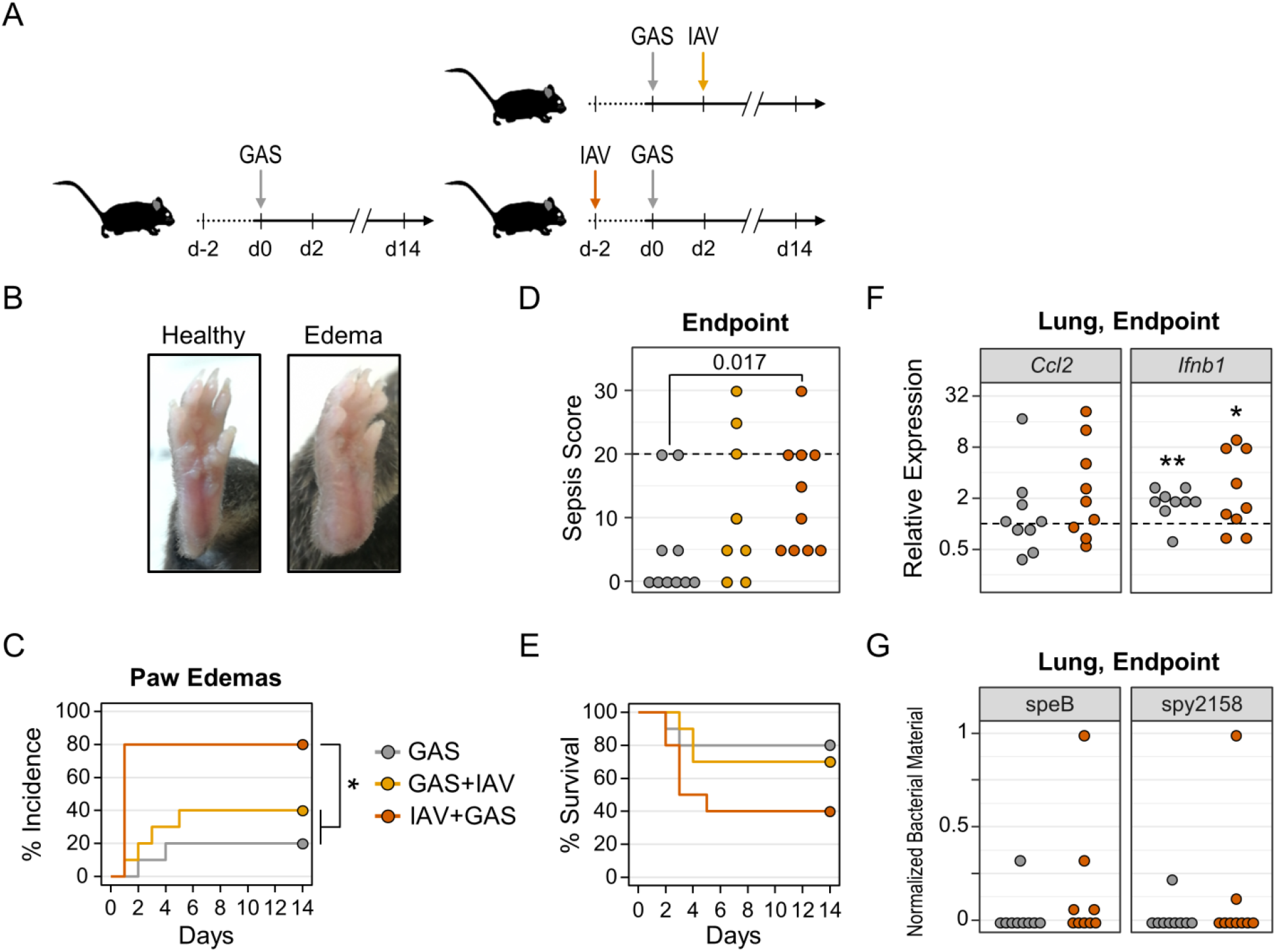
Preceding IAV infection promotes bacterial dissemination and sepsis severity during co-infection. (**A**) Experimental design. For monocausal bacterial infection, Group A Streptococcus was administered intravenously (GAS, n = 10). For co-infection, mice were either infected with GAS followed by intranasal IAV administration (GAS+IAV, n = 10) or infected with IAV followed by infection with GAS (IAV+GAS, n = 10). (**B**) Representative photographies of a healthy compared to an edematous paw after IAV+GAS co-infection. (**C**) Kaplan-Meier curves display the incidences of paw edemas. *p < 0.05, log-rank test with p-value adjustment for multiple comparisons (Bonferroni-Holm method). (**D**) Dotplot shows sepsis scores at endpoints (day 14 or humane endpoint). p-value results from Mann-Whitney U test. The dashed line indicates the minimum score for humane endpoints. (**E**) Kaplan-Meier curves display survival probabilities. (**F**) Dotplots show endpoint bulk lung mRNA gene expressions of *Ccl2* and *Ifnb1*that were normalized to *Gapdh* and lungs from uninfected mice (dashed line) by the 2^-ΔΔCt^ method. *p < 0.05, **p < 0.01, one-sample Wilcoxon signed-rank test (μ = 1). (**G**) Normalized bacterial loads based on Ct values for GAS specific genes in endpoint lung homogenates.

Additionally, more blood smears and knee joint capsule swabs were positive for β-hemolytic bacteria in co-infected mice from the IAV+GAS group (Table I), which suggested that preceding influenza promoted bacterial dissemination and invasion into synovial tissues. By assessing macroscopic signs of burden as a proxy for sepsis severity (see Materials and Methods), we found a significantly increased median disease score when comparing post-influenza bacteremia with monocausal GAS infection (Fig. 2D). Interestingly, when correlating sepsis scores with eicosanoids from paw homogenates, we found a significant relationship between the individual disease severity and the corresponding amounts of prostaglandins D_2_ and E_2_ as well as 5- and 12-Hydroxyeicosatetraenoic acid (supplementary Table III). Furthermore, elevated disease severity in the IAV+GAS group was paralleled by a reduction in survival probability to 40% compared to 80% in the GAS only group (Fig. 2E). In contrast, mice from the GAS+IAV group had an only marginally decreased survival chance of 70%. However, the overall probability for a fatal outcome was, according to logrank statistics, not significantly different between groups (p = 0.13) and this was likely due to low sample sizes and a high degree of uncertainty.

**Table I.**
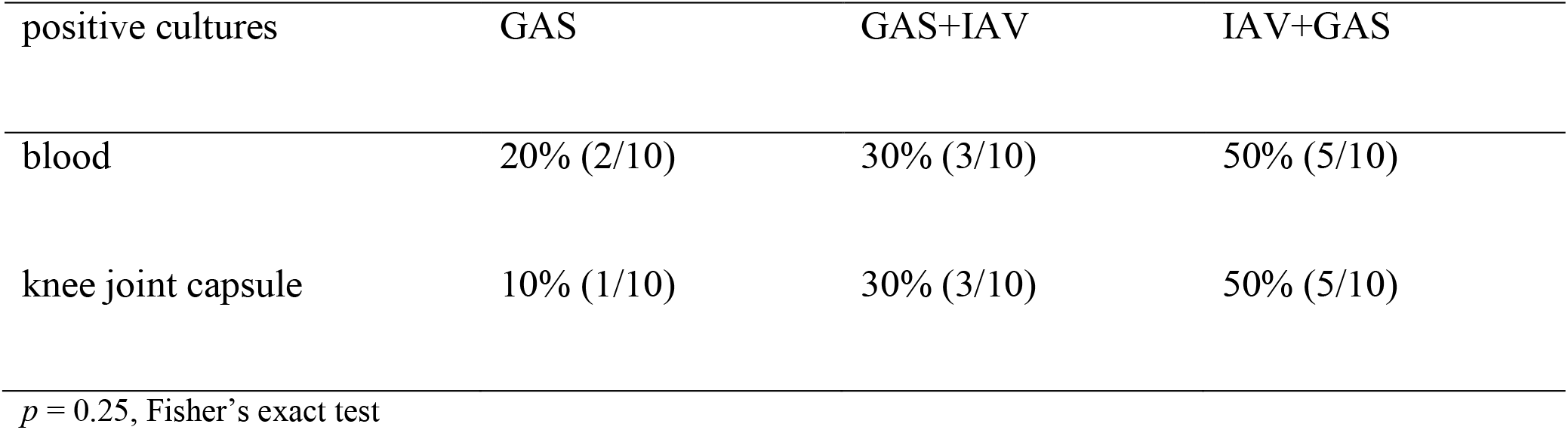
Frequencies of blood agar cultures from endpoint blood smears and synovial knee joint swabs positive for β-hemolytic bacteria.

| positive cultures | GAS | GAS+IAV | IAV+GAS |
| --- | --- | --- | --- |
| blood | 20% (2/10) | 30% (3/10) | 50% (5/10) |
| knee joint capsule | 10% (1/10) | 30% (3/10) | 50% (5/10) |

$p = 0.25$ , Fisher’s exact test

For our further analyses, we focused on the IAV+GAS co-infection sequence because our data suggested that the clinical outcome was not different between the GAS+IAV and GAS groups. We next aimed to investigate whether post-influenza GAS infection impacted on the immune activation in the lower respiratory tract. To this extent, we analyzed lung homogenates for the expression of *Ccl2* and *Ifnb1*, and compared the data from co-infected mice to GAS mono-infection or uninfected controls. We found that neither GAS nor IAV+GAS infection resulted in a meaningful alteration of the *Ccl2* expression in the lung (Fig. 2F, left panel). Of note, lungs from both mono- and co-infected mice exhibited a median 2-fold upregulation of *Ifnb1* relative to lungs from uninfected animals (p = 0.008 for GAS and p = 0.039 for IAV+GAS; Fig. 2F, right panel). Yet, when comparing the infection regimens with each other, we found that *Ifnb1* overexpression was comparable between both infection groups (p = 0.93).We were curious whether the bacteria are capable of disseminating from the blood into the lower respiratory tract and therefore analyzed lung homogenates for the presence of GAS specific genes using quantitative PCR (supplementary Fig. 5). Indeed, we detected genomic speB in four out of nine lungs from the IAV+GAS group whereas only one out of nine lungs from the GAS group was positive for this bacterial gene (Fig. 2G, left panel). However, when analyzing for spy2158, only two lung extracts from the IAV+GAS group were positive (Fig. 2G, right panel). Specific amplification was again confirmed by melting curves (supplementary Fig. 6). As whole lungs were submitted to chaotropic agent assisted homogenization and PCR analysis, we were not able to confirm whether there were any vital bacteria present in these samples.

Collectively, our *in vivo* data demonstrated that a preceding IAV infection of the respiratory tract aggravated intravenous GAS infection by promoting localized inflammation and a dysregulated host response as shown by an elevated sepsis scores. In contrast, application of the virus following an already established bacteremia did not influence on disease progression and outcome.

### 3.3 Preceding influenza A virus infection impacted on the Group A Streptococcus induced diversification of macrophage surface expression profiles

As our *in vivo* co-infection model implicated a preceding IAV infection to cause impaired control of the bacterial challenge following a superimposed GAS infection, we sought to explore any modification of anti-bacterial innate immunity. Macrophages are considered as first line defense immune cells that are substantial to contain pathogens in early phases of infection [43]. These cells take part in numerous bacterial infectious diseases and are especially crucial for a competent innate immune response during invasive GAS infection [31,44–46]. Hence, we chose *in vitro* (co-)infection models of primary murine macrophages in order to investigate whether IAV influences on the GAS-induced immune landscape. In detail, murine macrophages were differentiated from bone marrow cells by M-CSF stimulation and were subsequently infected with IAV, GAS or IAV and GAS (Fig. 3A). We then analyzed the expression patterns of immunologically relevant surface antigens by flow cytometry. In order to gain insight into differentially expressed macrophage markers, we performed dimension reductions on our multiparametric data sets by t-distributed stochastic neighbor embedding (t-SNE). Fig. 3B demonstrates for the topological distribution of surface marker expression levels distinct allocations of cells that were obtained from the different infection models. For instance, macrophage subsets overexpressing CD80 and CD86 were seemingly enriched in IAV+GAS co-infected cultures, whereas mono-infection with GAS resulted in the accumulation of CD206 overexpressing macrophages. Unsupervised clustering of macrophage populations on the basis of their respective expression patterns by flowSOM further indicated that co-infection triggered a different response than viral or bacterial mono-infections (supplementary Fig. 7).

**Figure 3.**
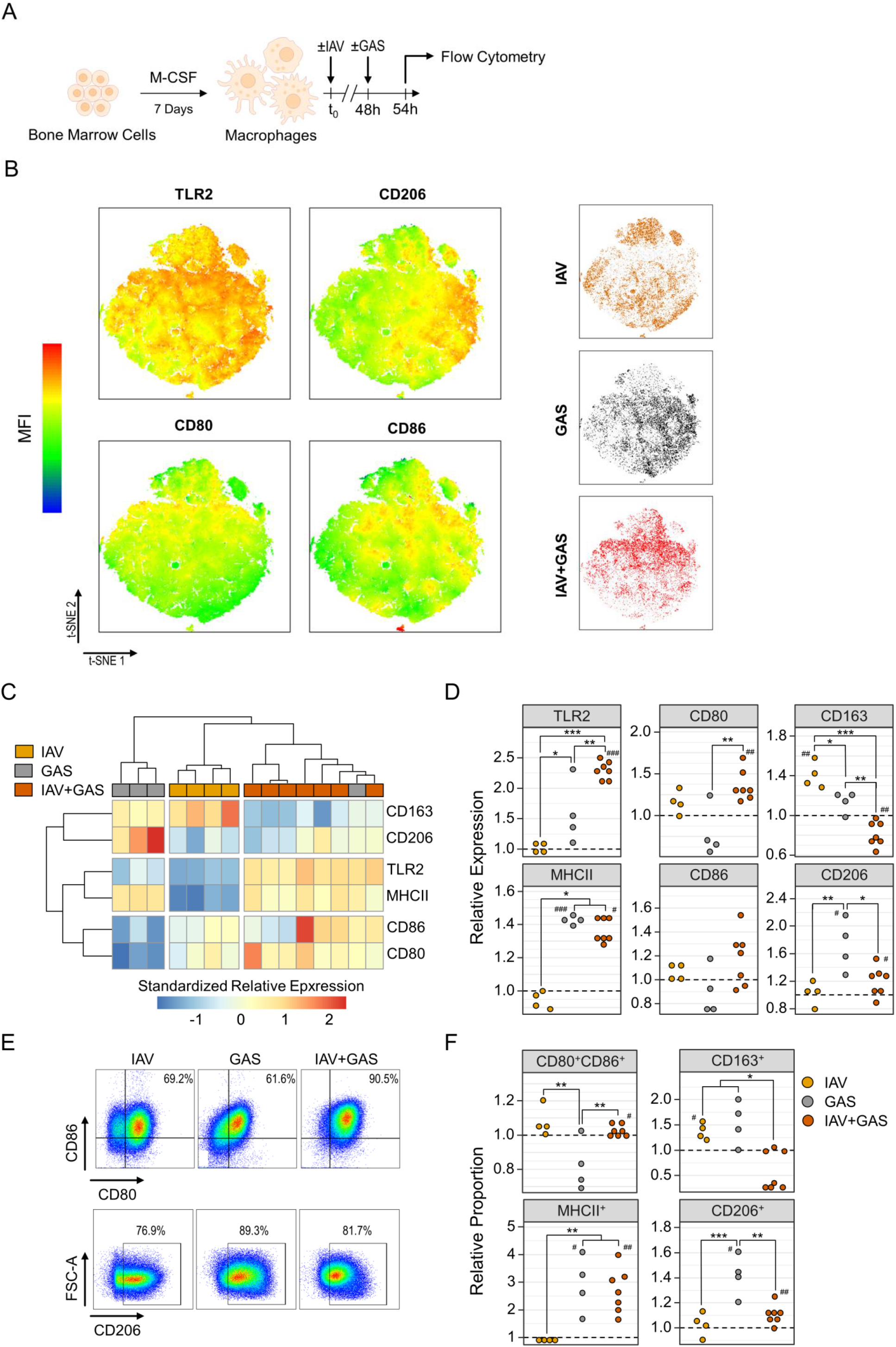
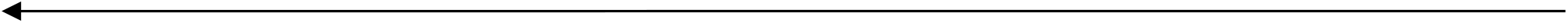
Activation of bone-marrow derived murine macrophages during *in vitro* co-infection was distinct from bacterial mono-infection. (**A**) Experimental Design. Macrophages were differentiated from bone marrow cells and then infected with either IAV for 48 h (n = 4) or GAS for 6 h (n = 4). For co-infection, IAV was first applied for 48 h followed by GAS infection for 6 h (IAV+GAS, n = 7). Each sample was obtained from individual mice to obtain biological replicates. (**B**) t-distributed stochastic neighbor embedding (tSNE) on flow cytometry data from the three different macrophage infection models and topology of surface antigen expression levels. 10,000 events from each sample were integrated into the dimension reduction analysis. MFI: median fluorescence intensity. (**C**) Heatmap and hierarchical clustering on standardized fold changes of surface antigen expression levels based on their MFI. Fold changes were generated by normalization of MFI data from infected macrophages to their respective paired uninfected controls. (**D**) Dotplots depict the alteration of surface antigen expression levels due to (co-)infection. (**E**) Representative pseudocolor plots illustrate the alteration of proportions of macrophages expressing CD80 and CD86 (top) or CD206 (bottom) after (co-)infection. (**F**) Dotplots demonstrate the shift of macrophage subpopulation fractions after (co-)infection relative to uninfected controls (dashed lines). *p < 0.05, **p < 0.01, ***p < 0.001, Dunn’s test or Tukey HSD test with p-value adjustments for multiple comparisons (Bonferroni-Holm method). ^#^p < 0.05, ^##^p < 0.01, ^###^p < 0.001, Wilcoxon signed-rank test or one-sample t-test for the comparison to uninfected cultures (μ = 1).

In an effort to obtain a more detailed picture of IAV- and GAS-induced immune responses, we next focused on the individual expressions of macrophage surface antigens. Given the inter-experimental variance of macrophage cultures, median fluorescence intensities (MFI) of (co)-infected cells were normalized to their corresponding uninfected controls that were acquired from the same donor animal (supplementary Fig. 8). Notably, expression patterns were similar within each group, which resulted in a robust hierarchical clustering for IAV, GAS and IAV+GAS infected macrophages (Fig. 3C). In detail, apart from a significant upregulation of CD163 compared to both bacterial infection and co-infection, IAV had hardly any impact on the expression of the investigated surface proteins (Fig. 3C, 3D). Conversely, GAS infection induced the over-expression of TLR2, which was even amplified following co-infection (Fig. 3D). Both the applications of GAS and IAV+GAS comparably prompted an elevated production of MHCII. Although not statistically significant, GAS infection led to a slight downregulation of CD80, which was reversed to an upregulated expression in the IAV+GAS group. Similarly, co-infection triggered a minor overexpression of CD86 that was short of reaching statistical significance due to a high within-group variance (p = 0.067, compared to uninfected). The downregulation of CD163 as well as the attenuation of the GAS-induced CD206 upregulation in the IAV+GAS group further supports the notion that a preceding IAV infection led to a distinct immune response in macrophages during co-infection (Fig. 3D).

As a result of differentially affected expression landscapes, the proportions of distinctive macrophage subpopulations shifted depending on the infection regimen (Fig. 3E). We found a minor depletion of CD80^+^CD86^+^ cells following GAS infection (p = 0.1), whereas co-infection caused a significantly increased proportion of this population when compared to uninfected controls (Fig. 3F). Both bacterial mono-infection and co-infection induced an enrichment of MHCII^+^ macrophages, suggesting a retained ability of these immune cells to inform and coordinate an adaptive immune response. In accordance with the altered expression profiles shown in Fig. 3D, the proportions of CD163^+^ and CD206^+^ cells, respectively, were decreased upon co-infection relative to GAS infection only (Fig. 3F).

Collectively, our data on the diversification of surface antigen expression demonstrated that the immune response of macrophages towards co-infection with IAV and GAS was considerably distinct from the effects that were induced by either mono-infection. Although we encountered some similarities between the GAS and IAV+GAS groups, the preceding viral infection seemingly manipulated or obliterated the macrophages’ reaction towards the bacterial pathogen.

### 3.4 A preceding influenza A virus infection impaired the inflammatory capacity of macrophages during co-infection

In order to gain further insights into the immune response triggered by infected macrophages, we next performed analyses on mRNA expression of immune mediators by quantitative PCR and measured cytokine secretion by ELISA or bead-based multiplex analysis (Fig. 4A). As illustrated in Fig. 4B, the different (co-)infection regimens triggered distinct expression patterns of immunomodulatory agents that resulted in strong within-group associations as shown via hierarchical clustering. IAV infection was specifically characterized by a relatively higher expression of *Mgl2* and *Tgfb1* (Fig 4C, supplementary Fig. 9). Bacterial infection, on the other hand, comprehensively stimulated the overexpression of several genes that mediate an inflammatory response (Fig. 4B). Strikingly, co-infected macrophages mostly failed to induce a similar magnitude of GAS inducible overexpression, yet upregulated *Arg1* (Fig. 4B, 4C).

**Figure 4.**
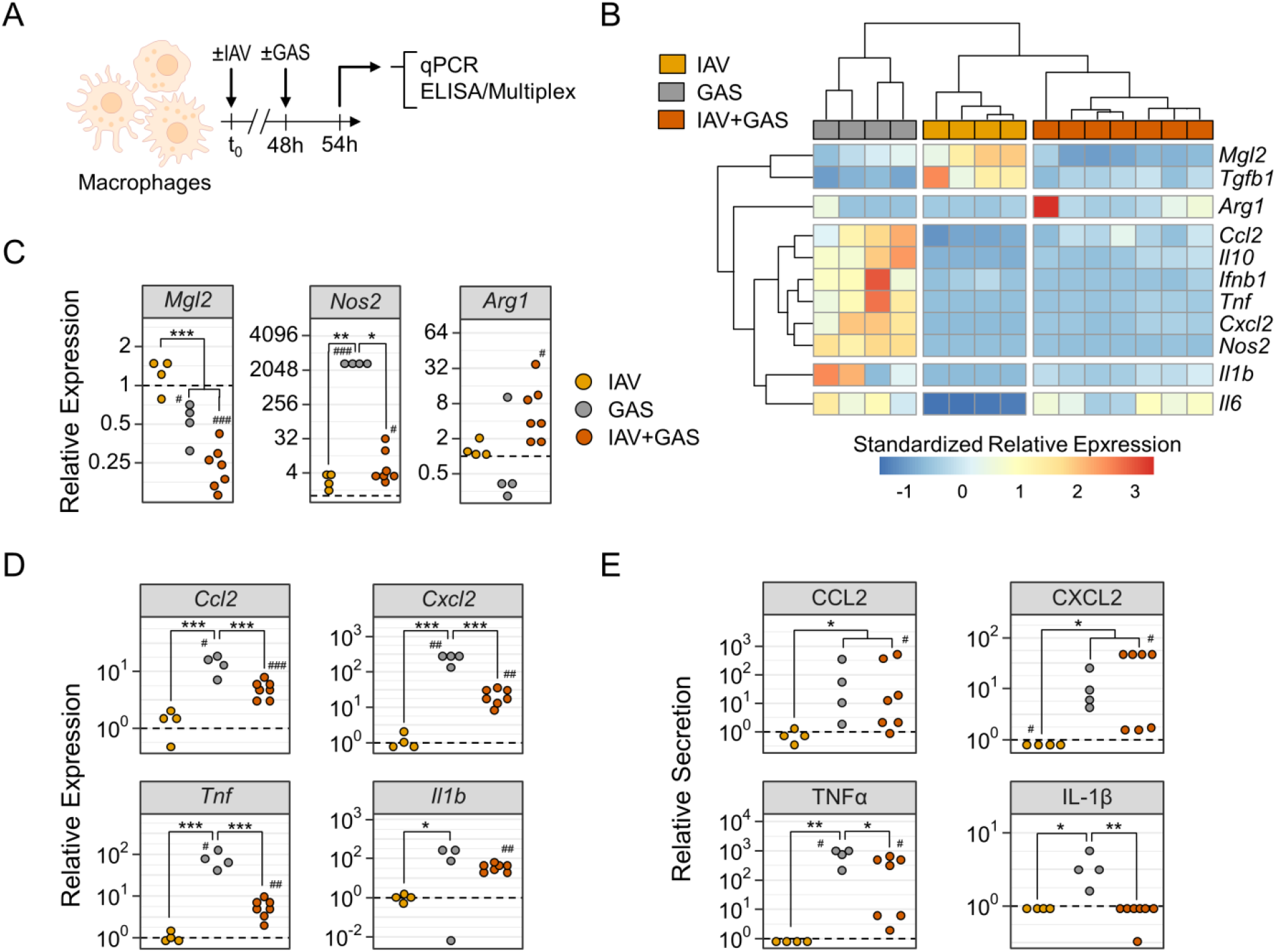
Preceeding influenza A virus infection impedes pro-inflammatory immunological features of macrophages during co-infection. (**A**) Experimental Design. Bone-marrow derived macrophages were infected with IAV (n = 4), GAS (n = 4) or IAV+GAS (n = 7). Each sample was obtained from individual mice to obtain biological replicates. (**B**) Heatmap and hierarchical clustering on standardized relative mRNA expression levels from quantitative PCR analyses using the 2^-ΔΔCt^ method. Data from infected cultures were normalized to *Gapdh* and their respective paired uninfected controls. (**C**) Dotplots show the alterations of *Mgl2, Nos2* and *Arg1* mRNA expression levels due to (co-)infection. (**D**) Dotplots illustrate distinct patterns of chemokine and cytokine mRNA production by macrophages after (co-)infection. (**E**) Dotplots demonstrate (co-)infection induced protein production of chemokines and cytokines that were measured in cell culture supernatants. Dashed lines represent control cultures. *p < 0.05, **p < 0.01, ***p < 0.001, Dunn’s test or Tukey HSD test with p-value adjustments for multiple comparisons (Bonferroni-Holm method). ^#^p < 0.05, ^##^p < 0.01, ^###^p < 0.001, Wilcoxon signed-rank test or one-sample t-test for the comparison to uninfected cultures (μ = 1).

By further examining individual expressions, we found that *Mgl2* was significantly reduced following bacterial mono-infection and co-infection by 1.8- and 3.9-fold, respectively, compared to uninfected controls (Fig. 4C). Remarkably, GAS application induced an approximately 3,000 fold overexpression of *Nos2* that was impeded during co-infection to a mere, yet statistically significant, 10-fold overexpression. Furthermore, co-infection triggered the upregulation of *Ccl2, Cxcl2* and *Tnf*, which were significantly less pronounced in comparison to GAS infection only (Fig. 4D). Secretion of these cytokines was mostly comparable between these groups, however TNFα production by co-infected macrophages was reduced (Fig. 4E). Although both *Il6* and *Il10* were increased in the GAS and IAV+GAS group, respectively, only co-infection caused a significant secretion of the protein products (supplementary Fig. 9). While *Ifnb1* was only upregulated following GAS infection, *Tgfb1* was downregulated after GAS infection as well as co-infection (supplementary Fig. 9A). Of note, although the GAS-induced overexpression of *Il1b* was also observed in the IAV+GAS group, co-infection entirely abrogated the secretion of mature IL-1β, which suggests that a preceding IAV infection compromised innate immune sensing of Streptococci (Fig. 4D, 4E).

In summary, the expression patterns of immunologically active mediators were noticeably different between GAS mono-infection and IAV+GAS co-infection, implying that prior virus infection modifies anti-streptococcal immunity.

## 4 Discussion

In this study we demonstrated that influenza promoted subsequent intravenous GAS infection and allowed for dissemination of the bacterial pathogen within the blood and its migration into lungs as well as synovial tissues. Although we did not assess any alterations in bone or cartilage morphology, we would like to argue that an invasion of articular tissue by GAS is reminiscent of septic arthritis [47]. Indeed, we previously demonstrated that the occurrence of paw edemas, which in the present study was more likely during IAV and GAS co-infection, was due to bacterial colonization of both, subcutaneous and periarticular tissues and was paralleled by immune cell infiltration [35]. Hence, we here show for the first time, that a preceding IAV infection predisposes the host to severe complications during GAS blood infection. Conversely, IAV infection elicited subsequent to intravenous GAS infection did not aggravate disease severity, suggesting that immune priming events in response to a prior viral encounter mitigate an otherwise competent anti-bacterial immune response.

Influenza in humans is usually characterized by mild-to-moderate disease that is rarely lethal and resolves shortly after infection [48], which was also shown in our animal model of IAV inoculation. Upon entry into nasopharyngeal cavities, the virus trespasses into the mucus, invades the epithelium and spreads to immune cells [49,50]. The host then recognizes parts of the viral RNA genome by intracellular pattern recognitions receptors, which triggers the production of several inflammatory cytokines, among them type I IFNs, that establish an anti-viral immune state [51–53]. We have demonstrated that residual viral genes persisted for 16 days in the lungs of some infected mice, which was paralleled by a continuous upregulation of *Ifnb1*. However, we believe it to be unlikely that replicative viral particles were still present in the lungs up to this point because IAV is typically cleared within a couple days following infection and the quantities of viral genes were barely detectable in our samples [54–56]. Type I IFN can have beneficial effects during bacterial infection by promoting host resilience and by preventing systemic hyperinflammation [57–60]. However, several studies advocated that the consequences of type I IFN expression are detrimental for the containment of a secondary bacterial insult subsequent to influenza [20,61,62].

By using a mouse strain that lacks the common IFNα/β receptor (IFNAR) in a model of pneumococcal superinfection, Shahangian and colleagues demonstrated that the IAV-induced IFNAR signaling led to an impaired production of the neutrophil attractants CXCL1 and CXCL2 [22]. They argued that, in agreement with a complementary study by Didierlaurent *et al*., type I IFNs desensitize subsequent TLR-mediated recognition of bacterial components by macrophages, which are major producers for these chemokines [22,23]. Another work on IFNAR^-/-^ mice by Nakamura and colleagues had some contrasting results concerning the impact of type I IFN signaling on pneumococcal superinfection [24]. In their study, they found that the virus and the bacteria were capable of synergistically inducing an overproduction of type I IFNs, which led to an impaired production of CCL2 while CXCL1/2 production was unaltered [24,63]. CCL2 supports bacterial clearance by the attraction of CCR2^+^ monocytes to the infected tissue [64,65]. Along these lines, we found in our study that CCL2 was significantly reduced in the plasma of IAV-infected mice and that both, monocausal bacterial infection and co-infection featured *Ifnb1* over-expression in the lung. Hence, although the role of CCL2 during GAS infections is not yet fully elucidated, we find it possible that a preceding influenza restricts anti-bacterial immunity by limiting monocyte homing and their differentiation to macrophages not only in pulmonary tissues but also in remote host compartments that would be affected during GAS blood infection. However, the cellular source of this chemokine was not identified in our study. Furthermore, the expression of *Ccl2* in lung samples that were taken at endpoints was comparable between bacterial infection and co-infection, which challenges the idea that IAV-induced alterations in immune cell recruitment continues after bacterial superinfection. In order to delineate the progression of co-infection in more detail, future studies should therefore focus on observations that are performed at specific time points rather than taking samples at endpoints that might be difficult to compare.

Apart from the ramifications due to an impaired chemokinogenesis, we suspected other means by which IAV dampens innate immune sensing of GAS. We hence focused on macrophage immunobiology in the context of co-infection and found that the virus comprehensively altered GAS-induced gene expression patterns and cytokine layout. In detail, we detected that the immune sensors CD163 and CD206 were markedly downregulated in co-infected compared to GAS only infected macrophages. CD163 is an acute phase-regulated scavenger receptor that is exclusively expressed by cells of the monocyte lineage and aids in the removal of potentially toxic iron complexes during intravascular hemolysis [66–69]. Due to the fact that CD163 also mediates tissue repair [70], host resilience [66,68], immune resolution and is able to sense gram-positive bacteria [71,72], we speculate that this receptor might confer a protective immune state during hemolytic bacteremia, even though its role in GAS infection is yet underexplored. Similarly, the mannose receptor CD206 might support pathogen sensing during co-infection [73–77], however mice that lack this sensor molecule are not more susceptible to infection [78,79].

Strikingly, a preceding IAV inoculation notably reduced the GAS-induced upregulation of *Nos2* while boosting *Arg1* expression. Both genes code for enzymes that compete for the substrate L-Arginine, yet induce opposed immune mechanisms [80–82]. While nitric oxide synthase 2 (NOS2) provides inflammatory and bactericidal metabolites [83–85], arginase (ARG1) supports tissue repair and immune resolution [83]. Thus, our data hint at a distortion of anti-bacterial processes due to a prior IAV infection. This is further corroborated by an inadequate sensing of the bacterial pathogen indicated by the reduced and abolished production of TNFα and IL-1β, respectively, which was similarly shown in a model of pneumococcal superinfection [19]. Interestingly, we detected for both, GAS mono-infection and superinfection an upregulation of *Il1b*, which suggests that the incapacity of co-infected macrophages to process and secrete IL-1β is due to a failure in the GAS-inducible activation of the NLRP3 inflammasome [86–89]. In fact, it was shown that different variants of IAV, including a 2009 pandemic strain, were capable of thwarting IL-1β maturation by interfering with NLRP3 inflammasome assembly [90–92], which is crucial for innate immune sensing and coordination [93]. An IAV-mediated nullification of IL-1β secretion would be of dramatic consequences during streptococcal superinfections. The absence of signaling via the IL-1 receptor (IL-1R) was in fact associated with an increased susceptibility to systemic GAS infection in both mice and humans [86,94,95]. Remarkably, rheumatoid arthritis patients that received the IL-1R antagonist Anakinra exhibited a roughly 330-fold increased rate of invasive GAS infections which included an elevated likelihood of life-threatening complications such as necrotizing fasciitis and sepsis [95].

Although our study yielded several findings that are of interest to the research about influenza and bacterial co-infections, it has some limitations. Apart from the fact that our observations remained purely phenomenological and no experiments on underlying mechanisms were performed, we were unable to link the gap between our *in vivo* and *in vitro* models. For instance, the IAV-induced reduction of CCL2 in mice was not observed in macrophages that were challenged with the virus. Furthermore, it would have been interesting to test whether IAV impedes the capacity of macrophages to phagocytize GAS. Nevertheless, the multitude of IAV-inducible alterations to macrophage immunobiology in the context of GAS superinfection was surprising to us and warrant further investigations.

In summary, we here describe in complementary *in vivo* and *in vitro* co-infection models that IAV infection thwarts anti-streptococcal innate immunity. This finding warrants further investigations on the mechanisms underlying this phenomenon that sets the stage for post-influenza superinfection. As an important side issue, our work underscores the importance of regular vaccinations against influenza in order to avert bacterial superinfection and prevent fatal invasive GAS complications [10,96–100].

## Acknowledgment

The authors would like to thank Wendy Bergman for her excellent support during the work for this study. We also thank Dirk Koczan and Ildiko Toth for their help with the transcription analyses. Moreover, we like to express our gratitude to Karin Gerber and Chantal von Hörsten for their participation in the animal breeding and care at the animal core facility. The authors sincerely give their thanks to all staff and students that took part in the scientific discussions and experimental conduct.

## Ko-Infekt Study Group

Karen Methling^2^, Michael Lalk^2^, Ulrike Blohm^3^, Alexander Schäfer^3^, Bernd Kreikemeyer^4^ (Affiliations are listed on the title page)

## Author Contributions

J.V. and B.M-H. designed the study. J.V., M.B., E.W., M.M., D.S., K.M., U.B. and A.S. designed and performed the experiments and collected the data. J.V., M.B. and D.S. performed the statistical analyses. J.V. wrote the first draft of the manuscript. All authors contributed to the article and approved the submitted version.

## Abbreviations Used in This Article

In order of appearance:

IAV: Influenza A Virus
GAS: Group A Streptococcus
HA: Hemagglutinin
IL: Interleukin
IFN: Interferon
MDCKII: Mardin-Darby Canine Kidney II (cells)
TCID50: Tissue Culture Infectious Dose 50
THB: Todd-Hewitt Broth
CFU: Colony-forming Units
20-HETE: 20-Hydroxyeicosatetraenoic acid
13-HODE: 13-Hydroxyoctadecadienoic acid
PGE_2_: Prostanglandin E_2_
AA: Arachidonic Acid
DMEM: Dulbecco’s Modified Eagle’s Medium
FCS: Fetal Calf Serum
M-CSF: Macrophage Colony-stimulating Factor
RB: Running Buffer
7-AAD: 7-Aminoactinomycin
MFI: Median Fluorescence Intensity
t-SNE: t-distributed stochastic neighbor embedding
CCL2/MCP-1: Chemokine CC-Motif Ligand 2/Monocyte Chemoattractant Protein-1
TNFα: Tumor Necrosis Factor α
CXCL2/MIP2-α: Chemokine CXC-Motif Ligand 2/Macrophage Inflammatory Protein 2-α
ELISA: Enzyme-linked Immunosorbent Assay
ANOVA: Analysis of Variance
r: Pearson Product-moment Correlation Coefficient
SEM: Standard Error of the Mean
IFNAR: Interferon-α/β Receptor
NOS2: Nitric Oxide Synthase 2
ARG1: Arginase
NLRP3: NLR Family Pyrin Domain Containing 3
IL-1R: Interleukin-1 Receptor

## Supplementary Material

Contents:

Supplementary Figure 1. Gating Strategy for the analysis of murine bone marrow derived macrophages.

Supplementary Figure 2. PCR Product lengths for influenza A Virus genes.

Supplementary Figure 3. PCR product melting curves for influenza A Virus genes.

Supplementary Figure 4. Eicosanoid production was not differentially regulated in paws from co-infected mice.

Supplementary Figure 5. PCR product lengths for Group A Streptococcus genes.

Supplementary Figure 6. PCR product melting curves for Group A Streptococcus genes.

Supplementary Figure 7. FlowSOM clutering on flow cytometry data from infected macrophages.

Supplementary Figure 8. Surface antigen expression changes on macrophages after infection and co-infection.

Supplementary Figure 9. Expression and secretion of cytokines induced by infection or co-infection.

Supplementary Table I. Primers for quantitative polymerase chain reaction of influenza A genes.

Supplementary Table II. Primers for quantitative polymerase chain reaction of Group A Streptococcus genes.

Supplementary Table III. Pearson correlation analyses of sepsis scores with paw eicosanoids.

