## Supplementary Information for "Influenza A Virus Exacerbates Group A Streptococcus Infection and Thwarts Anti-bacterial Inflammatory Responses in Murine Macrophages"

Johann Aleith *et al.*

### Supplementary Material

Contents:

Supplementary Figures 1 – 9

Supplementary Tables I – III

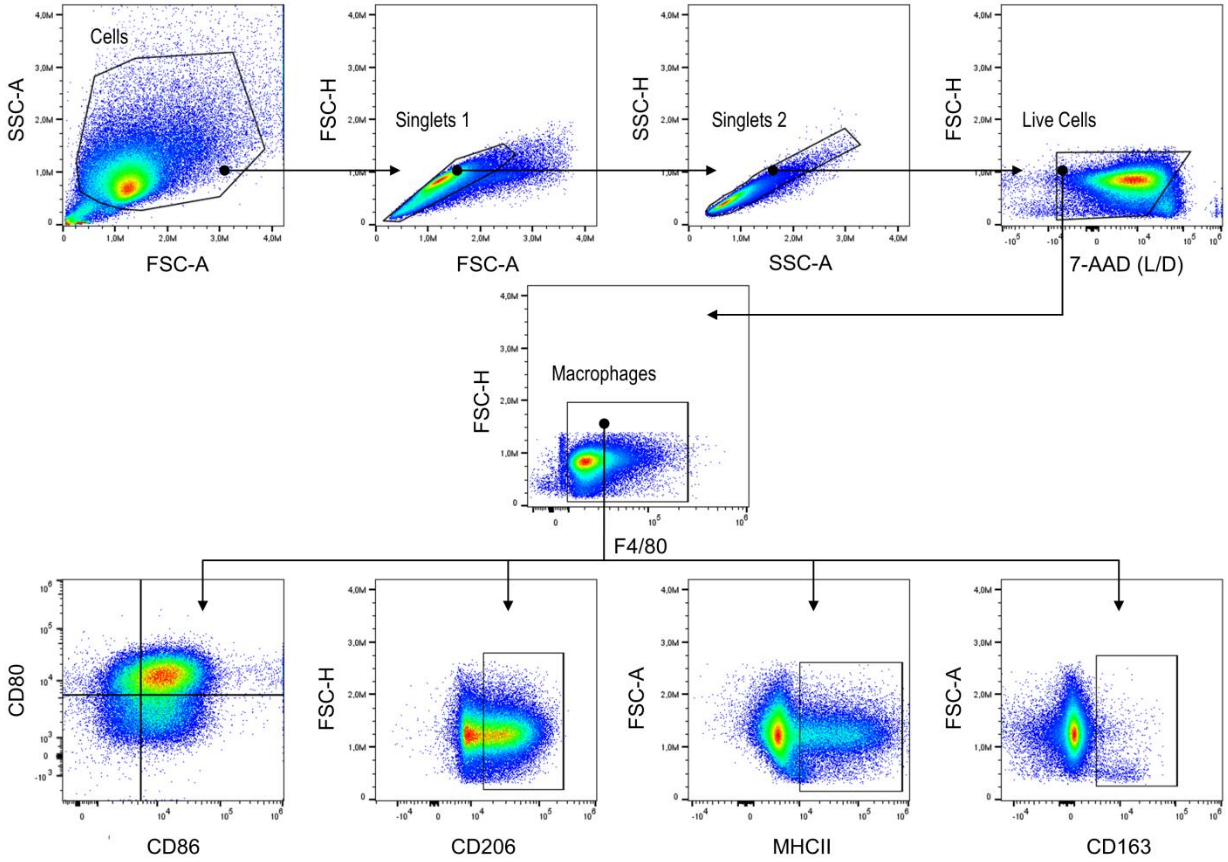


**Supplementary Figure 1. Gating Strategy for the analysis of murine bone marrow derived macrophages.** Debris and doublets were first excluded followed by the exclusion of dead cells that were positive for 7-AAD. Macrophages were identified by their expression of F4/80. Hierarchical gating was continued from this population and included the analyses of percentages of subpopulations that were $CD80^{+}CD86^{+}$, $CD206^{+}$, $MHCII^{+}$, and $CD163^{+}$, respectively.

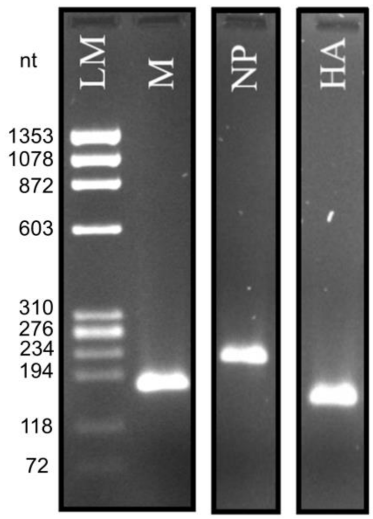


**Supplementary Figure 2. PCR product lengths for influenza A virus genes.** Primer pairs for the amplification of matrix protein (M), nucleoprotein (NP) and hemagglutinin (HA) were applied to reverse transcribed RNA extract from culture media that contained IAV. An agarose gel was run for quality control after PCR. Expected amplicon lengths were 179 nt for M, 236 nt for NP and 165 nt for HA.

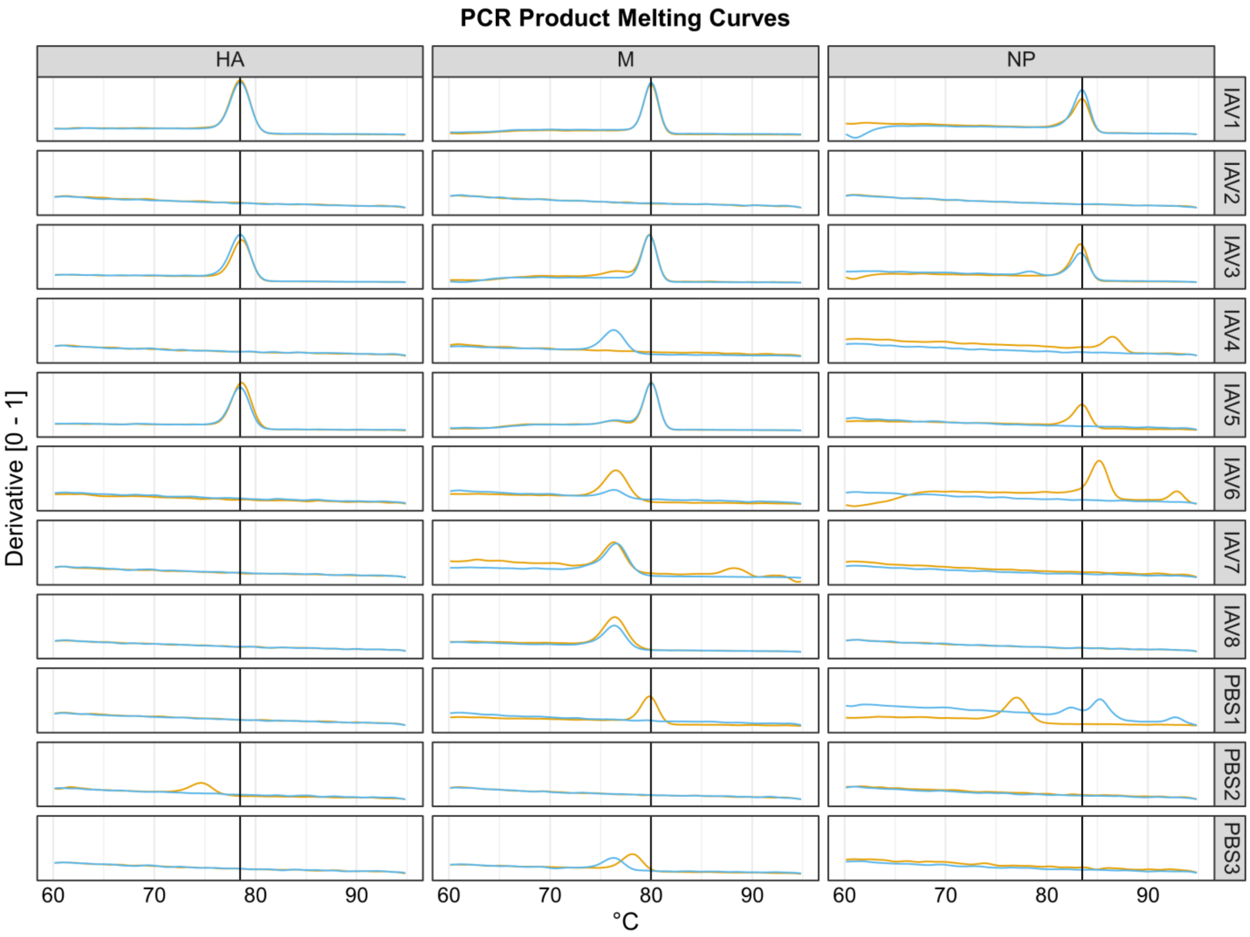


**Supplementary Figure 3. PCR product melting curves for influenza A virus genes.** Primer pairs for the amplification of matrix protein (M), nucleoprotein (NP) and hemagglutinin (HA) were applied to reverse transcribed RNA extract from culture media that contained IAV. Melting curves were determined after PCR. The derivates for the fluorescence signal changes are depicted. Vertical lines show the expected melting temperature of the amplicons which were 80°C for M, 83.5°C for NP and 78.5°C for HA, respectively.

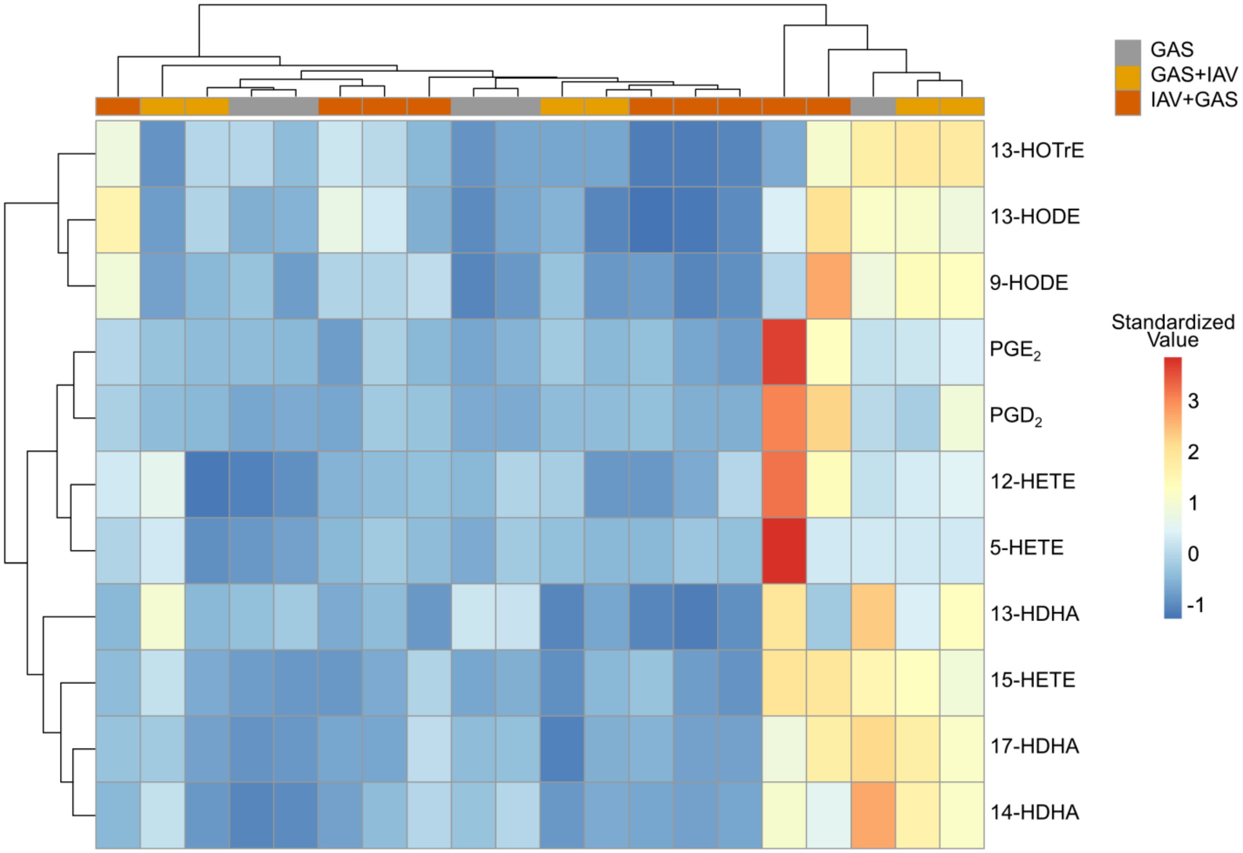


**Supplementary Figure 4. Eicosanoid production was not differentially regulated in paws from co-infected mice.** The heatmap shows the standardized amounts of eicosanoids that were extracted from mouse paws. Hierarchy doesn't show clustering between samples from the same infection group.

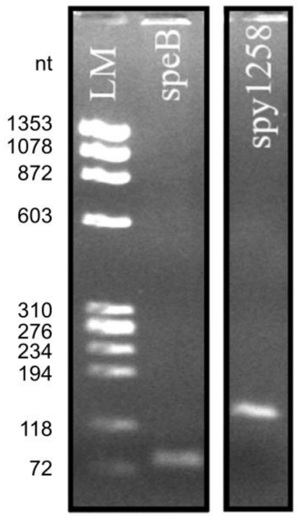


**Supplementary Figure 5. PCR product lengths for Group A Streptococcus genes.** Primer pairs for the amplification of streptopain (speB) and spy2158 were applied to DNA extracts from culture media containing Group A Streptococcus. An agarose gel was run for quality control after PCR. Expected amplicon lengths were 77 nt for SpeB and 136 nt for spy2158, respectively.

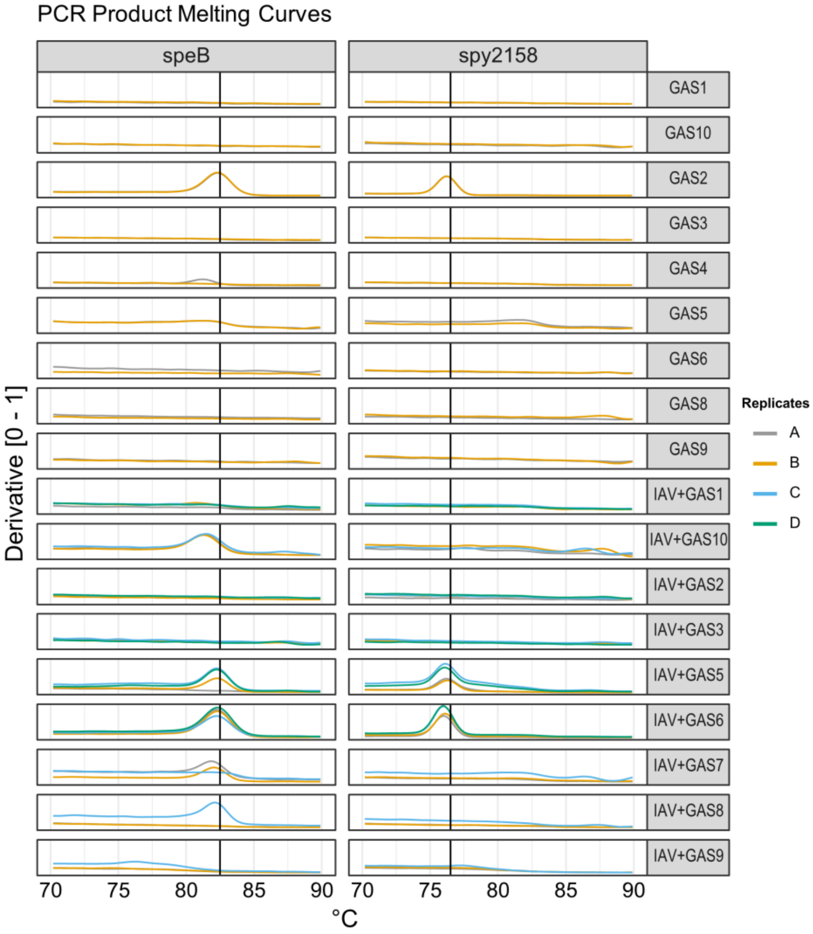


**Supplementary Figure 6. PCR product melting curves for Group A Streptococcus genes.** Primer pairs for the amplification of streptopain (speB) and spy2158 were applied to DNA extracts from culture media containing Group A Streptococcus. Melting curves were determined after PCR. The derivates for the fluorescence signal changes are depicted. Vertical lines shown the expected melting temperature of the amplicons which were 82.5°C for speB and 76.5 °C for spy2158, respectively. Twenty ng of DNA were used for PCR of technical replicates A and B. The maximum amount of DNA that was available was used for technical replicates C and D.

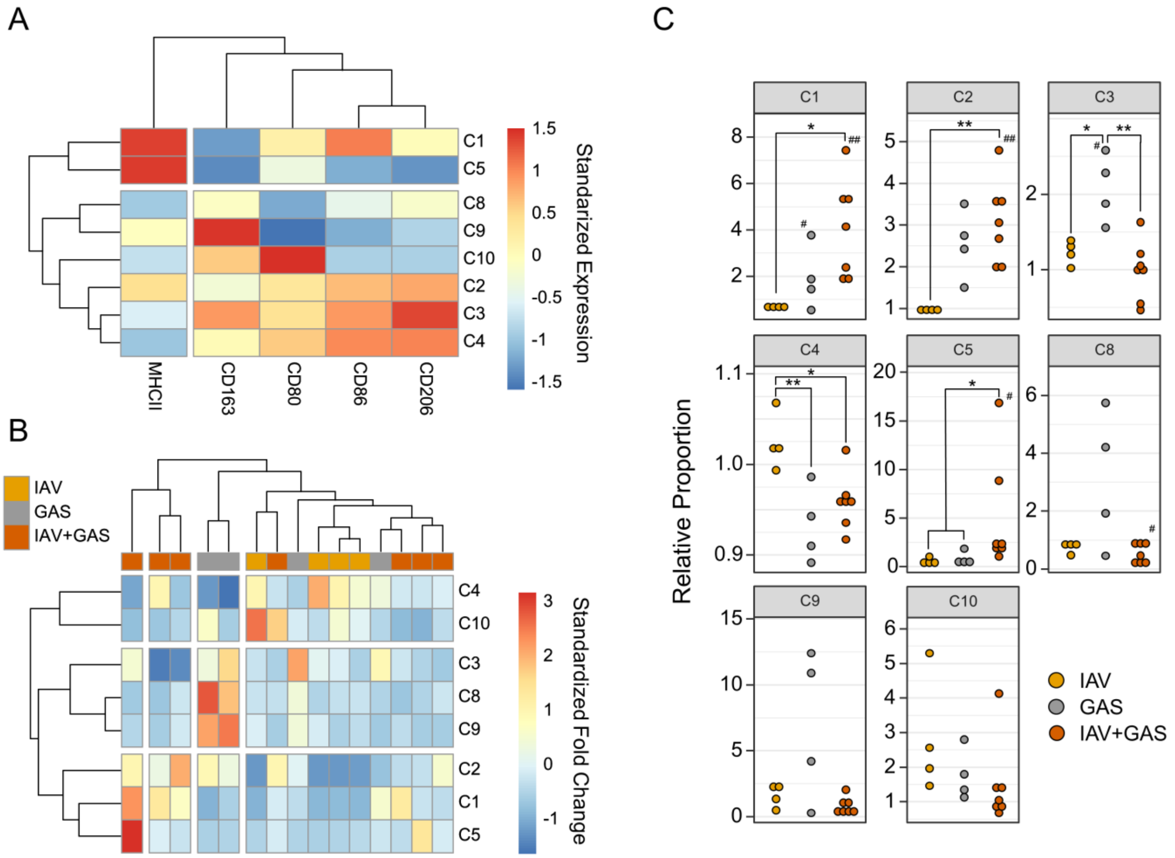


**Supplementary Figure 7. FlowSOM clutering on flow cytometry data from infected macrophages.** (**A**) Heatmap and hierarchical clustering on standardized surface antigen expression patterns that underlie the flowSOM-generated macrophage subpopulations (C1 – C10). C6 and C7 were excluded because they were not present in some samples. (**B**) Heatmap on standardized fold changes of flow-SOM generated macrophage subpopulation after infection with IAV or GAS or co-infection with IAV and GAS compared to the paired uninfected controls. (**C**) Dotplots showing alterations of flow-SOM generated macrophage subpopulations among infection groups relative to the paired uninfected controls. *$p < 0.05$, **$p < 0.01$, Dunn's test or Tukey HSD test with p-value adjustments for multiple comparisons (Bonferroni-Holm method). $^{\#}p < 0.05$, $^{\#\#}p < 0.01$, Wilcoxon signed-rank test or one-sample t-test for the comparison to uninfected cultures ($\mu = 1$).

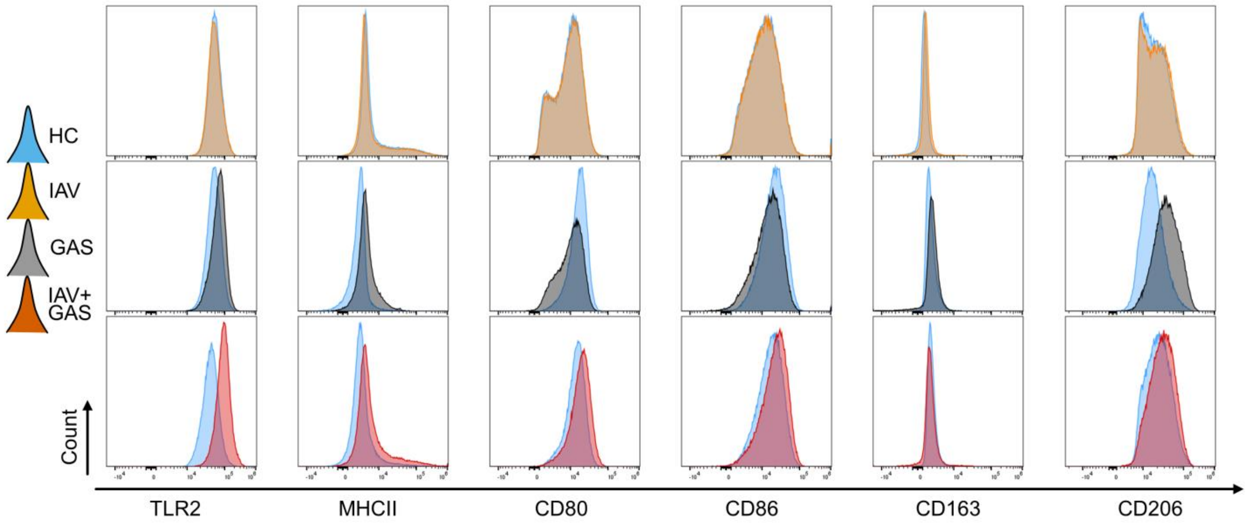


**Supplementary Figure 8. Surface antigen expression changes on macrophages after infection and co-infection.** Histograms depict the distribution of surface antigen expression of uninfected cultures (HC) and infected macrophages, respectively.

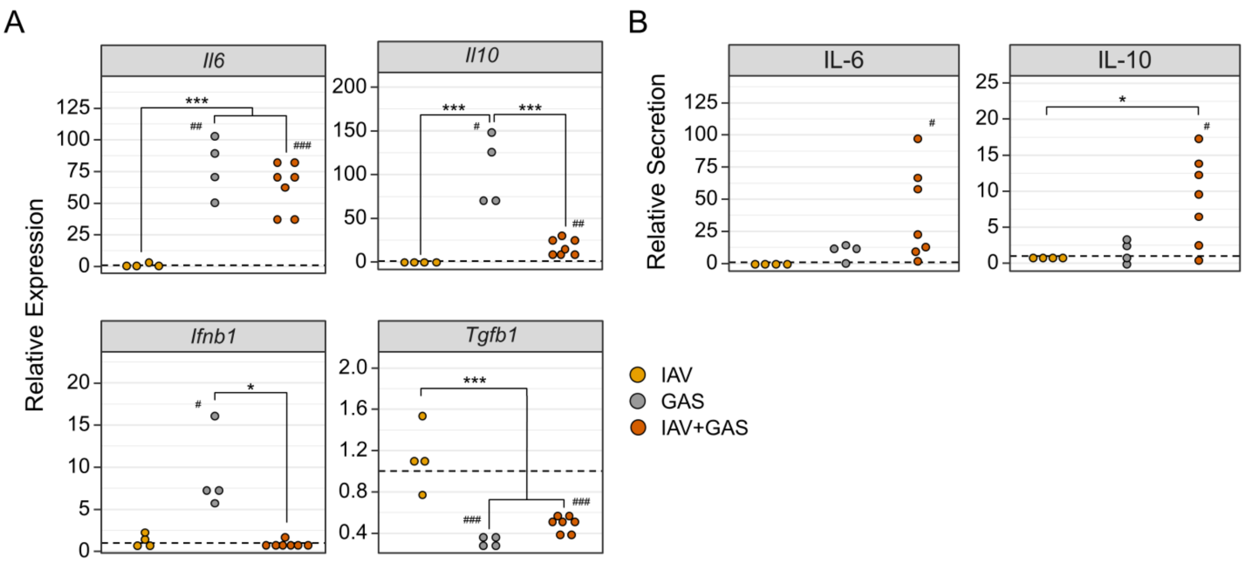


**Supplementary Figure 9. Expression and secretion of cytokines induced by infection or co-infection.** (**A**) Dotplots show relative mRNA expression data that were obtained by the $2^{-\Delta\Delta Ct}$ method. (**B**) Dotplots on cytokine data measured from supernatants of infected cultures relative to their uninfected paired samples. $^{*}p < 0.05$, $^{***}p < 0.001$, Dunn's test or Tukey HSD test with p-value adjustments for multiple comparisons (Bonferroni-Holm method). $^{\#}p < 0.05$, $^{\#\#}p < 0.01$, $^{\#\#\#}p < 0.001$, Wilcoxon signed-rank test or one-sampe t-test for the comparison to uninfected cultures (relative secretion = 1).

**Supplementary Table I.** Primers for quantitative polymerase chain reaction of Influenza A genes.

| Oligonucleotide Label | 5′ → 3′ Sequence | Amplicon Length | Melting Temperature |
|---|---|---|---|
| M fw | ACCAGAAGCGAATGGGAGTG | 179 nt | 80.0 °C |
| M rv | TCAGGCACTCCTTCCGTAGA | | |
| NP fw | TTCCACAAGAGGGGTCCAGA | 236 nt | 83.5 °C |
| NP rv | TCCGTCCTTCATTGTTCCCG | | |
| HA fw | GCCCGATCATGACTCGAACA | 165 nt | 78.5 °C |
| HA rv | CCCCATAGCACGAGGACTTC | | |

M: matric protein, NP: nucleoprotein, HA: hemagglutinin, fw: forward primer, rv: reverse primer

**Supplementary Table II.** Primers for quantitative polymerase chain reaction of Group A Streptococcus genes.

| Oligonucleotide Label | 5′ → 3′ Sequence | Amplicon Length | Melting Temperature |
|---|---|---|---|
| speB fw | CTAAACCCTTCAGCTCTTGGTACTG | 77 nt | 82.5 °C |
| speB rv | TTGATGCCTACAACAGCACTTTG | | |
| *spy2158 fw | ACCTCAAATTTCCGCAACTC | 136 nt | 76.5 °C |
| *spy2158 rv | TGCTCTCAATACTGGCAAGG | | |

fw: forward primer, rv: reverse primer, *primers were previously described by Dunne *et al.* (2013)

**Supplementary Table III**. Pearson correlation analyses of sepsis scores with paw eicosanoids.

| Eicosanoid | *r* | *p* |
|---|---|---|
| 12-HETE | **0.60** | **0.0053** |
| $PGD_2$ | **0.58** | **0.0079** |
| $PGE_2$ | **0.53** | **0.017** |
| 5-HETE | **0.52** | **0.02** |
| 15-HETE | 0.42 | 0.063 |
| 13-HDHA | 0.32 | 0.17 |
| 13-HODE | 0.30 | 0.2 |
| 9-HODE | 0.29 | 0.21 |
| 17-HDHA | 0.27 | 0.25 |
| 14-HDHA | 0.24 | 0.3 |
| 13-HOTrE | 0.15 | 0.52 |

r: correlation coefficient
